# Granger Sensori-Behavioral Taxonomy of Neuronal Ensemble Activity from Two-Photon Calcium Imaging Data

**DOI:** 10.64898/2026.05.12.724603

**Authors:** Sahar Khosravi, Nikolas A. Francis, Patrick O. Kanold, Behtash Babadi

## Abstract

Understanding how neuronal populations interact to encode and transform sensory information is a fundamental challenge in computational neuroscience. Most existing studies, however, study neural encoding, behavioral readout, and functional connectivity as disjoint problems. Two-photon calcium imaging enables simultaneous recording of large neuronal ensembles *in vivo*, driven by diverse stimuli and eliciting distinct behaviors. However, extracting directional functional connectivity metrics as well as encoding and readout properties of neurons from such data remains difficult due to indirect and noisy observations of spiking activity, slow temporal dynamics, and the latent interplay between external stimuli and endogenous neural processes.

Here, we introduce a unified conceptual and operational modeling and inference framework for *directly* extracting functional Granger causal (GC) effects between neurons, from external stimuli to neurons, and from neurons to behavior, from two-photon imaging data, in the sense of Granger. Inspired by the intersection information framework, we also identify neurons that encode features of sensory stimuli that inform behavioral readout. The resulting GC networks together with the taxonomy of functional sensori-behavioral relevance, which we call G-taxonomy, provides a powerful statistical analysis framework, enabled by the integration of several techniques including state-space modeling and inference, variational inference, and point processes.

We applied the proposed framework to simulated and experimentally-recorded two-photon imaging from the mouse auditory cortex (A1) during both passive listening and active tone discrimination. Our simulation studies reveal significant improvement of our proposed methodology over existing techniques. Analysis of experimental data from the mouse A1 identifies distinct groups of cells with diverse sensori-behavioral relevance, as well as changes in functional connectivity associated with correct vs. incorrect behavior. In summary, this work provides a principled and data-driven methodology for uncovering directional interactions among the neurons, sensory stimuli, and behavior, all within the same statistical framework, offering new insights into how distributed cortical populations transform sensory inputs into behaviorally relevant representations.

**Author Summary:** The brain processes sensory inputs through the coordinated activity of large networks of neurons and produces readouts that elicit behavior. Understanding how information flows and is processed through these networks is a central goal of neuroscience. In this study, we present a new computational framework that identifies directional interactions among neurons in an ensemble as well as from sensory stimuli to neurons and from neurons to behavior. Utilizing the Granger formalism to identify directional effects, as opposed to common correlational measures, our framework extracts said effects directly from two-photon calcium imaging data. We tested our proposed method on both simulated data and recordings from the auditory cortex of mice during passive listening and active tone discrimination tasks. Our method revealed diverse groups of neurons in the auditory cortex with distinct functional roles and relevance to sensori-behavioral integration. Our framework provides a new way to study the flow of information in the brain and can be broadly applied to uncover neural computations across sensory and cognitive systems.

## Introduction

It is widely accepted that the brain extracts features of sensory stimuli by generating distinct patterns of activity known as neural representations [1–5]. The process by which the sensory information is mapped to neural representations is often referred to as neural encoding. These neural representations are also read out by downstream cortical areas that mediate behavioral outcomes. This process is often referred to as behavioral readout or neural decoding [6–13]. Most existing studies of brain function consider neural encoding and behavioral readout as independent problems: they either focus on understanding how the brain encodes sensory information (e.g., receptive fields), or aim at predicting the behavior given the neural representations (e.g., choice probability models). As such, the interaction between the brain and behavior is obscured, which significantly limits our understanding of brain function. In order to tap into higher-level functional processes such as learning and attention, it is crucial to develop models designed to simultaneously account for both.

In recent work, a new framework known as Intersection Information (II) has been proposed, which aims at extracting the information in the neural representation that intersects with that relevant to behavior [14–16]. This framework uses partial information decomposition [17–19] to capture the mutual information between the behavioral choice and the pair of stimulus and neural activity, while subtracting the portion that is common between the neural activity and behavior independent of the stimulus. This framework has shown great promise in shedding light into the mechanisms by which neural encoding and behavioral readout intersect [20–24].

Despite the promising nature of this approach, it has so far been limited to small populations of neurons due to challenges in nonparametric estimation. In addition, the behavioral space has so far been limited to simple binary decision-making tasks [14]. Extending this framework to study sophisticated sensori-behavioral interactions in a scalable fashion requires addressing several challenges. First, inferring II from ensemble neural data requires nonparametric estimation of partial information measures and their bias correction, which scales poorly with the network size due to its combinatorial nature. Second, each neuron is considered in isolation, which undermines the network-level effects relevant to behavior. Third, the relevant features of neural activity are required to be fixed *a priori* (e.g., average spiking rate).

Apart from the foregoing challenges, in application to data modalities that indirectly measure neuronal spiking, such as two-photon calcium imaging, the spike trains are not readily available for computing information measures. In the case of two-photon imaging, the fluorescence observations are noisy and temporally blurred surrogates of spiking activity. Existing approaches carry out the inference of information measures in a two-stage fashion: first, infer spikes using a deconvolution technique, and then compute firing rates and evaluate information measures [20–22]. These two-stage estimates are highly sensitive to the accuracy of spike deconvolution, and require high temporal resolution and signal-to-noise ratios [25, 26]. Furthermore, these deconvolution techniques are biased toward obtaining accurate first-order statistics (i.e. spike timings) via spatiotemporal priors, which may be detrimental to recovering higher-order statistics that are crucial for computing information measures.

We propose to cast this problem within the so-called Granger formalism [27–29], in order to address these shortcomings. Our approach is first and foremost inspired by the elegant machinery of II and partial information decomposition [14–16], by the known equivalences between Granger causality (GC) and directed information inference in certain statistical models [28, 30–32], and finally by the fact that the parametric nature of GC inference allows its integration with multivariate point process models [33, 34]. The essence of GC can be understood by considering two processes, *X*_*t*_ and *Y*_*t*_, and asking whether knowledge of process *X*_*t*_ significantly enhances our ability to predict *Y*_*t*+1_. If it does, a GC link from *X* to *Y* is established. This notion of directed connectivity has been widely used in the analysis of neuroimaging data as well as spiking neuronal ensembles [27, 33–36].

Utilizing several recent theoretical and algorithmic advances for GC inference [34, 35, 37] as well as direct network estimation from two-photon imaging data [38, 39], we propose a *Granger Sensori-Behavioral Taxonomy*, namely G-taxonomy, to characterize the relevance of neuronal activity to stimulus, behavior, and their intersection. The key idea is to include the stimulus and behavior within the network of *N* neurons and consider an augmented network of size *N* + 2. Then, the notion of GC link between two neurons can be extended to GC links from stimulus to neurons (i.e., stimulus encoding), from the neurons to behavior (i.e., behavior decoding), and the interaction of stimulus and behavior conditioned on neuronal activity (i.e., intersection). As such, we will label the neurons in an ensemble as **G**ranger **S**timulus encoding (GS), **G**ranger **B**ehavior decoding (GB), **G**ranger **I**ntersection (GI), or none of the above.

The key technical challenge in establishing the G-taxonomy is devising a precise statistical framework, with controlled false discovery rate, that takes noisy and blurred two-photon recordings from a limited number of trials and outputs the aforementioned labeling. To achieve this, we propose a dynamic Bayesian network model that integrates point processes [33, 40] with multivariate autoregressive models [29] to jointly capture the effect of stimuli on neuronal activity as well as behavioral responses, directly from two-photon fluorescence data without requiring intermediate spike deconvolution. Model inference is efficiently carried out through an integration of methods such as Pólya-Gamma augmentation, Iteratively Reweighted Least Squares (IRLS), Fixed Interval Smoothing, Kalman Filtering, and Expectation-Maximization [38, 41–46]. Finally, a hypothesis testing framework using the deviance difference statistics is used to quantify each detected effect via the Youden’s J-statistic [34, 47], followed by the Benjamini-Yekutielli procedure [48] to control the false discovery rate in ascribing the foregoing G-taxonomy to a neuronal ensemble.

We demonstrate the utility of our proposed G-taxonomy using carefully-designed simulated datasets as well as experimentally-recorded two-photon imaging data from the mouse auditory cortex during passive stimulus presentation and tone discrimination tasks. Our simulation results corroborate the robustness of the proposed statistical testing framework in revealing the GC links between neurons, arising from non-stimulus driven activity, as well as identifying GS, GB, and GI neurons. Application to experimental data reveals the utility of the proposed G-taxonomy in identifying neurons with distinct contributions to correct vs. incorrect behavioral outcomes, in terms of sensory encoding, behavioral readout, their intersection, or none. While consistent with existing results, this new taxonomy provides a unified, scalable, and statistically principled framework for bridging sensory encoding and behavioral readout and for probing how directed neuronal interactions support task performance.

In summary, the main contributions of our work include: **1)** leveraging the Granger formalism to assess the sensori-behavioral relevance of neurons within a *network*, as opposed to doing so in *isolation*, thanks to advances in scalable GC inference; **2)** learning the relevant features of spiking activity in a data-driven fashion via efficient parameter estimation, as opposed to restricting them *a priori*; and **3)** precise characterization of the statistical confidence in ascribing the GS, GB, and GI labels via J-statistics.

## Results

In this section, we demonstrate the utility of our proposed estimation framework through both simulation and experimental studies of neuronal activity in the mouse auditory cortex. We begin with a controlled simulation to validate recovery of directed links and neuron classes under known network structure. We then apply the framework to two experimental datasets from mouse auditory cortex: one recorded during spontaneous and stimulus-driven activity, and one during active tone discrimination behavior. Together these analyses test the framework’s ability to detect stimulus and choice-related interactions under increasingly realistic conditions.

### Overview of the Proposed Granger Sensori-Behavioral Taxonomy

Consider a canonical experimental setting in which an external stimuli, denoted by **s**_*t,l*_, is chosen at trial *l* and presented independently for a total of *L* trials, while the spiking activity of a population of *N* neurons are indirectly measured using two-photon calcium fluorescence imaging. Fig. 1 (forward arrow) shows the generative model that is used to quantify this procedure. The fluorescence observation in the *l*th trial from the *i* neuron at time frame *t*, denoted by 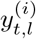, is a noisy surrogate of the intracellular calcium concentrations. The calcium concentrations in turn are temporally blurred surrogates of the underlying spiking activity 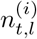, as shown in Fig. 1.

**Fig 1.**
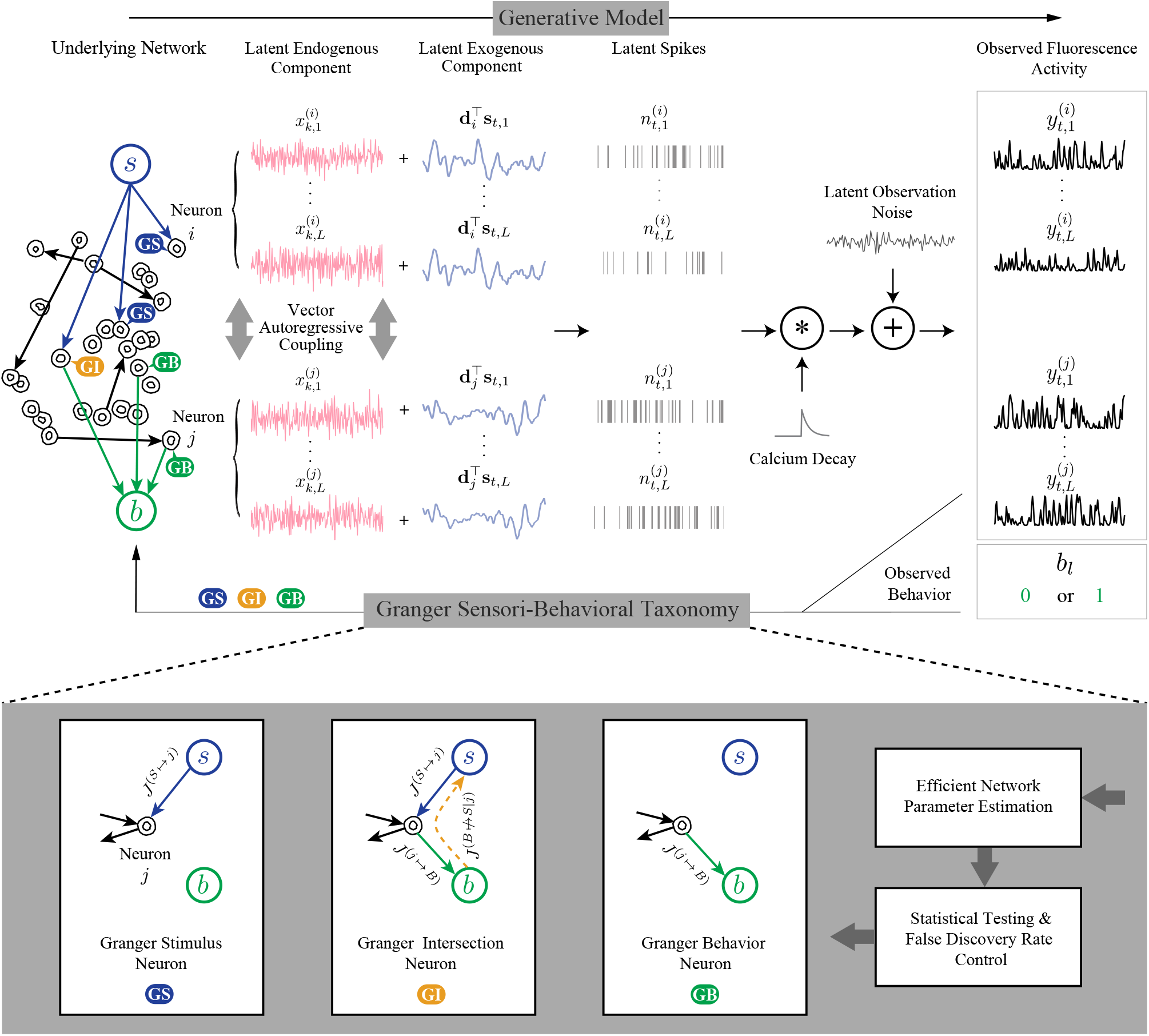
Schematic depiction of the generative model and proposed Granger sensori-behavioral taxonomy. Top: In the proposed model for estimating GC links directly from two-photon calcium fluorescence observations, observed variables (black), latent endogenous processes (red), and an exogenous stimulus (blue) relevant to the *i*th and *j*th neurons are shown. The spiking activity of networked neurons is influenced by latent endogenous processes (red traces) and an exogenous stimulus (blue traces), with black arrows indicating GC connectivity between neurons. The blue and green arrows, respectively, show the effects of the stimulus on neurons and neurons on behavior, in the sense of Granger. Bottom: our proposed inverse solution and G-taxonomy; following network parameter estimation, statistical testing, and false discovery rate control, we define three categories of neurons, namely Granger Stimulus, Granger Behavior, and Granger Intersection.

In modeling the spiking activity, we consider two main contributions: (1) the common known stimulus **s**_*t,l*_ affects the activity of the *i*th neuron via an unknown kernel **d**_*i*_, akin to the receptive field. We refer to this effect as the latent *exogenous* component; (2) the trial-to-trial variability and other intrinsic/extrinsic neural covariates that are not time-locked to the stimulus **s**_*t,l*_ are captured by a trial-dependent latent process 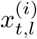. We refer to this effect as the latent *endogenous* component. Then, we use a Generalized Linear Model to link these underlying neural covariates to spiking activity [49].

To model the temporal coupling between the neurons, we assume that the *N*-dimensional latent endogenous process 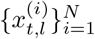 is governed by a vector autoregressive (VAR) model of order *P* with unknown parameters and process noise covariance. The binary behavioral outcome *b*_*l*_ at trial *l* is also modeled using a GLM, in which the neuronal activity within a “readout window” during or following stimulus presentation forms the set of relevant covariates.

Given this generative model, the objective of the inverse problem in Fig. 1 (backward arrow) is two-fold: using the observed data 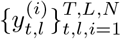 and 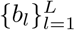, we seek to simultaneously (1) estimate the functional connectivity between the neurons in the sense of Granger, often referred to as Granger causality (GC), and (2) attribute a G-taxonomy to all the *N* neurons, in the sense of Granger, to quantify their role in the network function. The advantage of solving these two inverse problems simultaneously is also two-fold: (1) in detecting the GC links between neurons, the effect of external stimuli as a confound would be controlled. For instance, if neurons *i* and *j* are being driven by the stimulus with different latencies, by explicitly modeling the receptive fields **d**_*i*_ and **d**_*j*_ and quantifying the stimulus effect, the detected GC links would only be resulting from the VAR coupling of the endogenous latent process; (2) in attributing the G-taxonomy to each neuron, the network effects arising from endogenous interactions between the neurons as a confound would be ruled out. For instance, we can rule out if heightened activity of neuron *j* corresponding to *b*_*l*_ = 1 behavioral category is due to having a relevant effect to behavior, or is just due to receiving an endogenous coupling from a neuron *i* that directly contributes to behavior.

The proposed G-taxonomy is inspired by elegant machinery of intersection information and partial information decomposition [14–16], but is here carried out in the sense of Granger: here, an effect between two nodes in the network (including sensory stimulus and behavior), namely *source* and *target*, is quantified by assessing whether temporal forecasting of the target variable can be significantly improved by including the source variable and its time lags as covariates. Whereas in the partial information decomposition, an effect between two nodes is quantified by assessing whether the mutual information between their activity is significantly above chance level or not.

Using the Granger formalism, we perform network parameter estimation, followed by statistical testing and false discovery rate control to quantify the foregoing effects. Each effect or “link” is then statistically quantified by the Youden’s J-statistic, a popular summary statistic that includes both type-I and type-II error rates [34, 35]. That is, a J-statistic of 0 indicates a non-significant effect, and a J-statistic close to 1 implies a significant effect with small type-I and type-II error probabilities. As such, we can define three categories of neurons (Fig. 1, bottom panel):

- **G**ranger **S**timulus (GS) Neuron: as shown in the leftmost panel, neuron *j* is labeled as a GS neuron if there is a significant effect from the stimulus node to neuron *i*, i.e., *J*^*S* ↦ *j*^ is non-zero.
- **G**ranger **B**evahior (GB) Neuron: as shown in the rightmost panel, neuron *j* is labeled as a GB neuron if is exhibits a significant effect to the behavior, i.e., *J*^*j* ↦ *B*^ is non-zero.
- **G**ranger **I**ntersection (GI) Neuron: as shown in the middle panel, neuron *j* is labeled as a GI neuron if it is both GS and GB, and in addition in regressing the stimulus to the activity of neuron *j* during readout, including the behavior as an additional regressor does not significantly improve the regression accuracy, i.e., *J*^(*B* ↛ *S*|*j*)^ := 1 − *J*^(*B* ↦*S*|*j*)^ is non-zero.

Note that by virtue of the last definition, a neuron that is both GS and GB is *not* necessarily a GI neuron, as it may encode parts of the sensory information that are *not* informing the behavior, yet its activity could contribute to behavior. This is in fact the main rationale for defining intersection information [15]: the shared information between the behavior and the pair of *{*stimulus, neuronal activity*}* may contain a portion that pertains to features of the stimulus that are encoded in neuronal activity but do not inform behavior. Through a refinement of the partial information decomposition framework, this irrelevant portion is removed from the shared information to form II [15].

Here, we control for this irrelevant portion through quantifying *J*^(*B* ↛ *S*|*j*)^: if a neuron is encoding features of the stimulus that inform behavior, then regressing the stimulus to the neuronal activity during the readout window would not benefit from including the behavior as an additional covariate. Conversely, if a neuron is encoding features of the the stimulus that *do not* inform behavior, yet its activity during the readout window affects behavior, then its activity during the readout window is only a poor regressor of the stimulus, and including the behavior as an additional covariate is expected to improve the regression accuracy.

By a careful combination of *J*^*S* ↦ *j*^, *J*^*j* ↦ *B*^, and *J*^(*B*^ ↛ ^*S*|*j*)^, we will account for both type-I and type-II errors in attributing a GI label to a neuron. Finally, a neuron that fails to fall into any of these categories will remain unlabeled. The resulting G-taxonomy forms a novel labeling of the neurons in the network, as shown in Fig. 1 (top left network schematic).

### An Illustrative Simulation Study

To highlight the advantages and performance of the proposed G-taxonomy, we consider an illustrative simulation setting as shown in Fig. 2-A corresponding to a detection task. The network consists of *N* = 10 neurons observed over *T* = 10 s, with frame interval of 25 ms. The activity of the network is simulated for a total of *L* = 50 repeated trials, driven by a combination of exogenous stimuli and an endogenous VAR process of order *P* = 2, which is assumed to vary slower than the frame rate and hence piecewise-constant over *W* = 20 frames.

**Fig 2.**
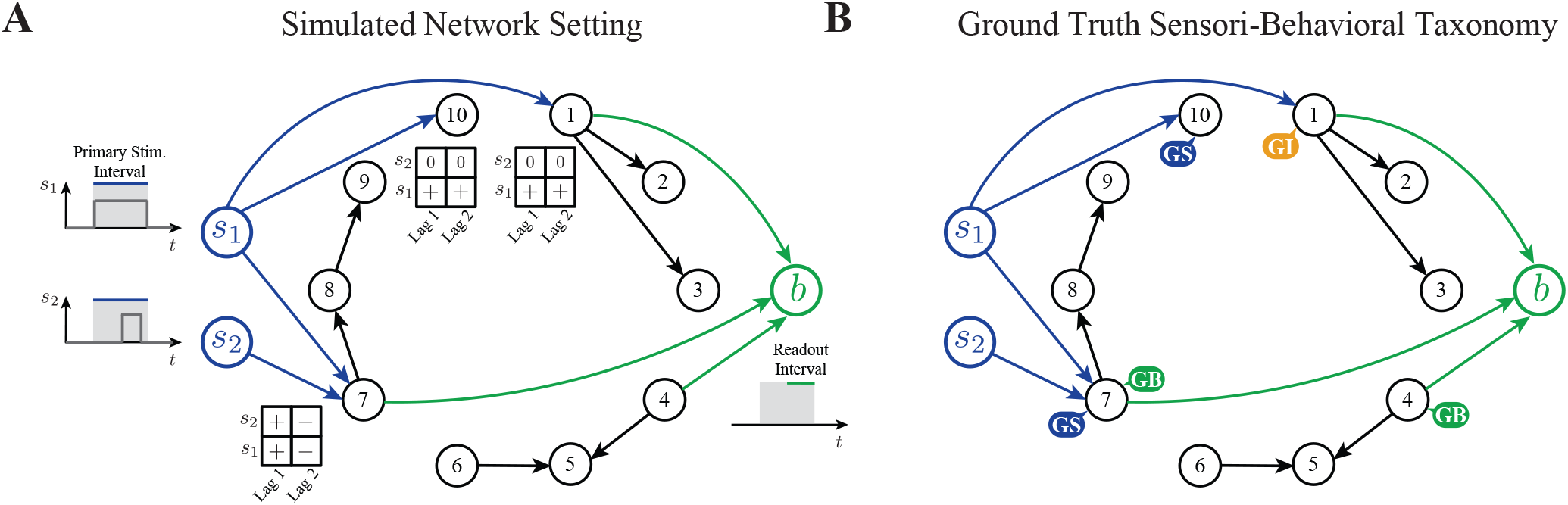
Results of the illustrative simulation study. A) Example of a simulated 10-neuron network. The time courses of the primary and secondary stimuli, as well as the behavioral readout window are shown as inset plots. The receptive fields of neurons 1, 7, and 10, which are influenced by the stimuli, are shown as 2-by-2 matrices. B) The ground truth Granger sensori-behavioral taxonomy of the network in panel A.

In each trial, the *primary* stimulus, *s*_1_, was randomly and uniformly turned “on” or “off”. In the “on” condition, the primary stimulus was a 100 ms pulse of magnitude 1. A background zero-mean white noise with variance 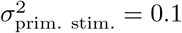 was added to the primary stimulus. In order to model a possible confounding effect, a *secondary* stimulus, *s*_2_, was also considered, which was in the form of a short pulse (quarter of the primary stimulus duration) occurring during the second-half of the primary stimulus interval, with the same magnitude and background noise variance (See the schematic time courses of *s*_1_ and *s*_2_ in Fig. 2-A). While the primary stimulus is chosen by the experimenter (e.g., intended visual or auditory stimulus), the secondary stimulus is a sensory stimulus that is not accounted for by the experimenter (e.g., unintended visual or auditory distractors). Nevertheless, both can be encoded by the neuronal network and affect the behavioral outcome [15].

Fig. 3-A shows two sample trials with the primary stimulus on (left) and off (right) for a sample neuron. Note that even when the primary stimulus is off, there could be significant spiking activity due to the fluctuations of the latent endogenous process (red trace in the first row). The binary behavior corresponding to the primary stimulus detection task was generated via a GLM model with covariates from from the average activity of a subset of stimulus-driven neurons (i.e., green arrows outgoing from neurons 1, 4 and 7 in Fig. 2) during the *readout interval*, defined as the second half of the primary stimulus interval (see the readout interval highlighted in Fig. 2-A). In the schematic network of Fig. 2-A, only neurons 1, 7, and 10 have a non-zero receptive field (i.e., blue arrows incoming from the stimuli). The receptive fields of neurons 1 and 10 are the same, and account for a sliding window average of the primary stimulus (i.e., both lags positive), whereas they are not responsive to the secondary stimulus. Neuron 7, however, has the same receptive fields for both stimuli, which model an onset/offset detector (i.e., the two lags have different polarities).

**Fig 3.**
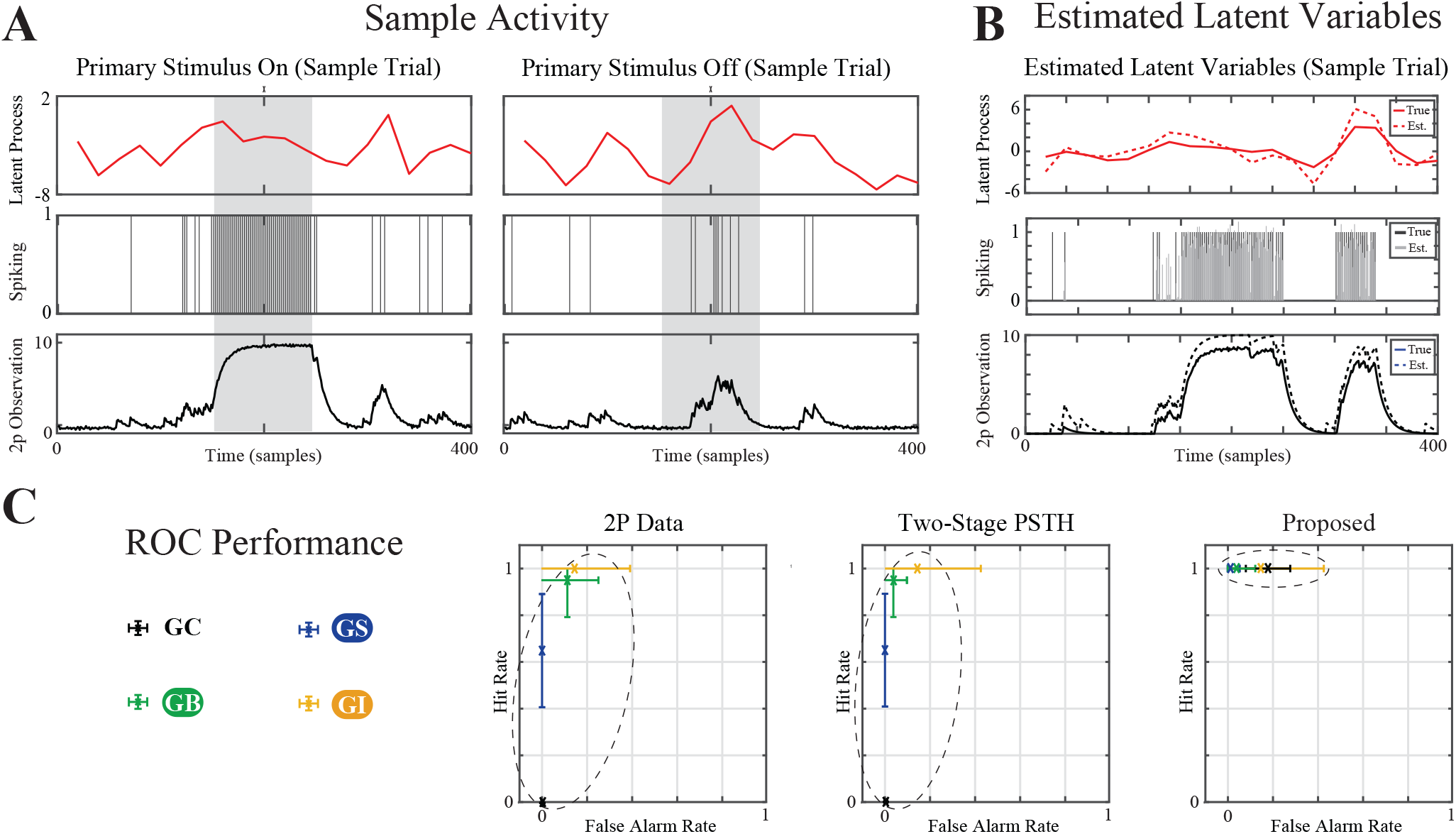
Results of the illustrative simulation study (cont’). A) Sample simulated trials with the primary stimulus on (left) and off (right). Rows (top to bottom) show the latent AR process (red), spiking activity (black), and resulting two-photon calcium traces (black). The shaded area show the primary stimulus interval. B) Sample trial showing estimated and ground-truth signals for a sample neuron (left). Rows top to bottom show estimated vs. true latent AR process, spiking activity, and calcium observations, respectively. C) The ROC performance of detecting the GC links between the neurons and the proposed G-taxonomy are shown for the 2P Data (left), Two-Stage PSTH (middle), and the proposed (right) methods. ROC analysis shows high sensitivity and specificity for all categories (FDR controlled) for our proposed method, whereas the 2p Data and Two-Stage PSTH suffer from poor hit rate and high false alarm rate (see the dashed ellipsoids for a visual guideline).

These receptive fields are schematically shown as 2-by-2 tables in Fig. 2-A.

The ground truth sensori-behavioral role of each neuron is shown in Fig. 2-B: neurons 1, 7, and 10 are GS neurons (i.e., non-zero receptive fields), and neurons 1, 4 and 7 are GB neurons (i.e., affecting behavior) by design. Neuron 1 encodes a moving average version of the primary stimulus, which is then read out during the readout interval and contributes to the behavior. As such, neuron 1 encodes stimulus information that informs behavior and is thus a GI neuron. Neuron 7, however, encodes the onset/offset of the primary stimulus *s*_1_, which does not significantly contribute to its activity during the readout window. Hence, it is encoding features of the primary stimulus that do not inform behavioral readout. As such, neuron 7 is both GS and GB, but not GI.

We expect the inferred G-taxonomy, obtained from the observed neuronal activity and behavioral outcomes only, to closely match the ground truth labels, which are determined by design. To generate various instances of such networks shown in Fig. 2-A, we constructed randomly generated networks by choosing three of the ten neurons at random to receive input from the primary stimulus, where one of them at random also received input from the secondary stimulus. The VAR coupling coefficients were kept the same across network realizations. Finally, in addition to the neuron receiving input from both stimuli, another stimulus-encoding neuron and a non-stimulus encoding neuron (chosen at random) were considered to form covariates for generating the behavioral outcome. Fig. 3-B shows an instance of the estimated endogenous latent process, spiking activity, and reconstructed observations for a sample neuron from a sample network. The estimated variables (dashed traces and gray bars) closely match the ground truth (solid traces and black bars).

Finally, Fig. Fig. 3-C shows the ROC performance curve for the proposed G-taxonomy over 10 network realizations, as well as those corresponding to two benchmark methods: *1) Two-stage PSTH:* in which the spikes are deconvolved first, using the FCSS procedure [43] to form the peristimulus time histogram (PSTH) of each neuron, followed by fitting a VAR model on said PSTH signals to obtain the network parameters and identify the taxonomy; *2) 2P Data*: in which the 2P data are considered to be governed by a VAR process, from which the network parameters and taxonomy are identified [20].

The proposed framework achieves perfect detection for the three categories as well as GC links between neurons. The false alarms were negligible for GS, GB, and GC categories, while they were slightly higher for the GI category. This is consistent with the difficulty of identifying the parameters of a GLM model with many parameters from a limited number of trials. The two-stage PSTH and 2P Data methods, however, exhibit poor hit rate and high false alarms, primarily for identifying the GC links and GS neurons. The details of statistical testing and FDR correction are given in the Methods.

In summary, our simulation study highlights how the proposed G-taxonomy can reveal distinct roles of neurons within a network: while specific neurons act as sensory encoding or behavioral readout neurons, there are some integrative nodes in the network that both encode sensory input and drive task-related behavior. These GI neurons bridge the encoding and decoding domains, that are often studied separately, and take the role of functional intermediaries in sensory–behavioral transformation.

### Identifying GS Neurons in Mouse A1 During Passive Listening

We next apply our proposed network inference and G-taxonomy to experimentally-recorded two-photon calcium imaging data from layer 2/3 of the mouse primary auditory cortex (A1). The dataset, originally reported by [38], consists of trials in which short broad-band noise stimuli are presented to the animal (referred to as *Stim. On* trials), which are randomly interleaved with silent trials of equal duration (referred to as *Stim. Off* trials). The stimuli were 75 dB SPL 100 ms broadband noise (4 − 48 kHz). Each trial was 5.1 s long (1 s pre-stimulus silence + 0.1 s stimulus + 3 s post-stimulus silence), and the inter-trial duration was 3 s. The recordings therefore contained both stimulus-driven and spontaneous neural activity, providing an opportunity to examine both GC links between neurons as well as their stimulus encoding properties.

We analyzed a subset of *N* = 9 neurons selected at random, with *L* = 50 stimulus presentation trials per neuron and *T* = 135 frames (5.1 s) per trial. The fluorescence traces (Δ*F/F*) of each neuron were normalized based on the average peak amplitude of prominent transients, ensuring consistent scaling across cells. The observation noise covariance matrix was estimated from the first 30 frames of each trial, corresponding to pre-stimulus baseline activity.

To capture the temporal receptive field structure of each neuron, we considered 10 lags of the stimulus, representing responses over a 330 ms window. The effective integration window for the latent AR process was set to *W* = 10 frames, assuming that neural activity within this window was driven by the same latent cortical process. The target FDR was chosen as 0.01.

The estimated network is shown in Fig. 4-A, with the detected GS neurons labeled.

**Fig 4.**
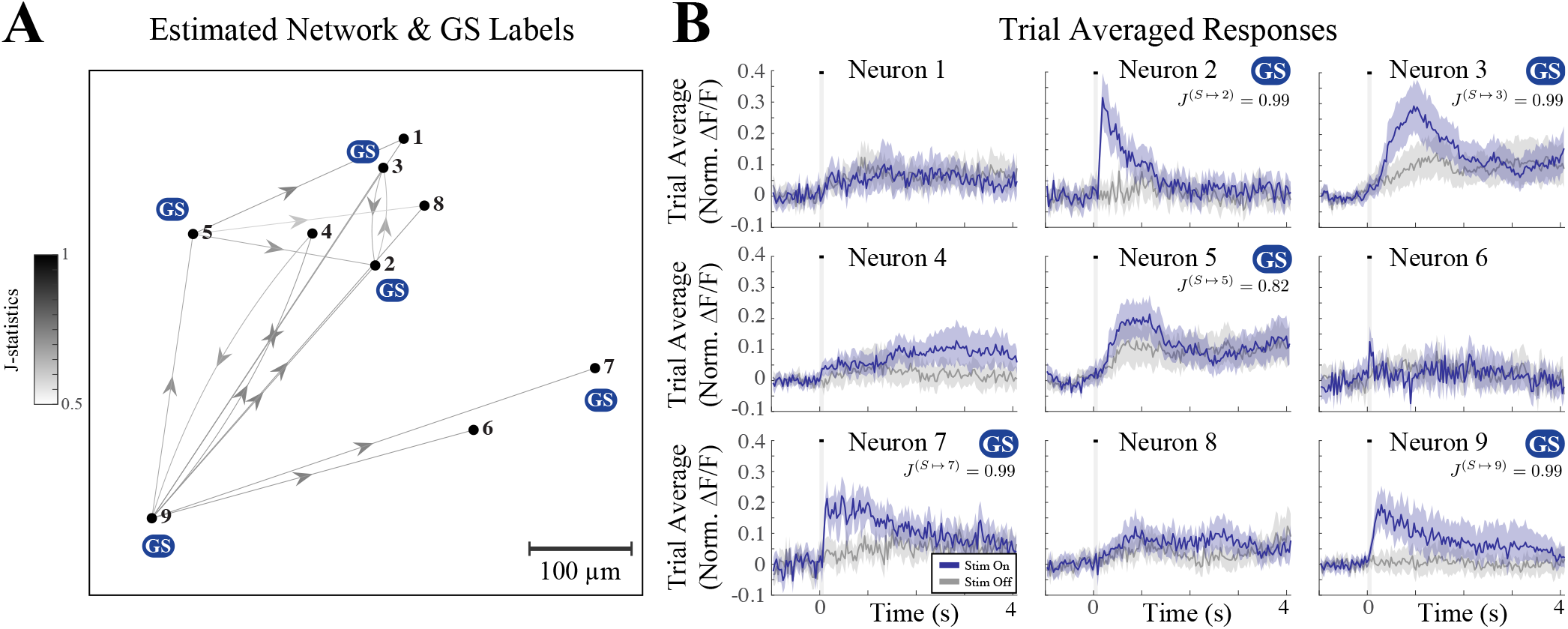
Identifying GS neurons in mouse A1 during passive listening. A) Estimated GC network and GS labels in mouse A1 during passive listening to randomly interleaved trials of silence and white-noise stimuli. B) The trial averaged responses of the neurons in panel A for Stim. Off (gray) and Stim. On (blue) trials. The stimulus interval is shown as a gray vertical hull. The J-statistics of the GS neurons are shown as insets in each sub-panel. The colored hulls show 95% confidence intervals.

While some of the GS neurons also participate in the GC network as sources, e.g., neurons 5 and 9, others are not (e.g., neurons 1 and 7). To further validate the detected GS labels, Fig. 4-A shows the trial averaged responses of all neurons for Stim. Off (gray traces) and Stim. On (blue traces) trials. As it can be visually confirmed, the responses of the GS neurons in Stim. Off and On trials are quite distinct, where are they are widely overlapping for the non-GS neurons. The J-statistics are also reported in the insets, which are consistent with these visual observations: the J-statistics for neurons 2, 3, 7, and 9 with visibly distinct response profiles under both trial conditions are 0.99, which shows high confidence in labeling them as GS. Neuron 5, however, is detected as GS with a J-statistic of 0.82, which is consistent with the response profiles being more similar.

Our analysis reveals two key advantage of the proposed G-taxonomy: first, the GS labels are detected conditioned on the entire network activity, which can thereby rule out potential network effects that could be otherwise attributed to stimulus. For instance, the response profiles of neuron 4 under Stim. Off and On trials are dissimilar later throughout the trial, and a statistical test performed on their difference may conclude that this neuron is responsive to the stimulus. However, based on our proposed analysis, this observed difference is likely due to the GC link from another GS neuron, i.e., neuron 9, which exhibits similar response profiles later throughout trial, while being highly responsive to the stimulus onset.

Second, the parametric nature of the underlying statistical tests in detecting GC links as well as the G-taxonomy are likely to have higher statistical power than commonly-used non-parametric tests applied to response differences shown in Fig. 4-B. This is particularly important when the number of trials and the trial duration are limited. For instance, while neuron 5 is detected as a GC neuron, a simple non-parametric statistical test applied to the response differences would not register as significant. This situation becomes even more stringent after correcting for multiple comparisons across neurons. However, our method tests for the improvement in response prediction variance by including the stimulus as a covariate on a trial to trial basis, and thereby labels neuron 5 as a GS neuron with a J-statistic of 0.82. This implies that while the type I error (i.e., p-value) is less than 0.01 due to the choice of the FDR rate, the test power is at least 0.82.

In summary, our analysis shows that the proposed network inference and G-taxonomy can be used as an alternative to existing approaches in detecting significantly responsive neurons under passive listening to randomly interleaved silent and stimulus-driven trials, with the advantage of ruling out possible network effects as well as increasing the test power compared to commonly-used non-parametric tests for responsiveness.

### Granger Sensori-Behavioral Taxonomy of Neurons in Mouse A1 L2/3 During Tone Discrimination

To test how our framework works in behaving animals, we analyzed two-photon calcium imaging data from the mouse A1 L2/3 during a pure-tone discrimination task [50]. Head-fixed mice performed a go/no-go task while A1 L2/3 activity was recorded. Low-frequency tones (7 or 9.9 kHz) were targets (go) and high-frequency tones (14 or 19.8 kHz) were non-targets (no-go); all four frequencies were randomly interleaved. The animals were trained to wait 0.5 s after the stimulus onset before making a decision (lick or no lick). Trials were labeled as hit (H), miss (M), correct rejection (C) or false alarm (F) based on the first lick [50]. We analyzed both passive blocks, where tones were presented while the mouse was quiescent, and behavior blocks with active task performance with the same tones. Detailed experimental procedures (animal training, imaging, and behavioral definitions) are provided in [50].

Unless otherwise noted, our analyses used a subset of 10 experiments from the data in [50]. The 10 experiments contained 1798 total neurons in the field of views, 225 of which exhibited significant activity in response to tones. Within each selected experiment, we randomly sampled *N* = 10 neurons and used all *L* = 50 available trials with *T* = 135 imaging frames per trial. For each neuron, Δ*F/F* traces were normalized by the average amplitude of visually identified peak transients to place cells on a comparable scale. The observation noise covariance was estimated from the first 30 frames of each trial (pre-stimulus baseline) and pooled across trials to form a per-neuron variance estimate. Stimuli were represented as binary pulse trains (tone onsets) and we modeled the receptive fields with with two temporal lags. The effective integration window for the latent AR process was set to *W* = 10 frames (330 ms), assuming that neural activity within this window was driven by the same latent cortical process.

Fig. 5–A summarizes the distribution of detected links across passive and behavior blocks (top) as well as the pooled trial averaged response of all neurons in correct (H & C) vs. incorrect (F & M) trials (bottom). The number of detected links during the passive blocks are significantly larger than that during incorrect trials. In addition, the number of links during incorrect trials is significantly smaller than that for both behavior categories pooled together. These findings are consistent with the results of [20] using short integration windows to analyze the connectivity changes during passive and behavior conditions. The average responses in correct (black) and incorrect (red) trials shown in the bottom panel exhibit intervals of significant difference both during the peri-stimulus and post-stimulus intervals. The time points of significant difference are marked by small vertical bars below the graph, with a family-wise error rate of less than 5% (See Methods for details).

**Fig 5.**
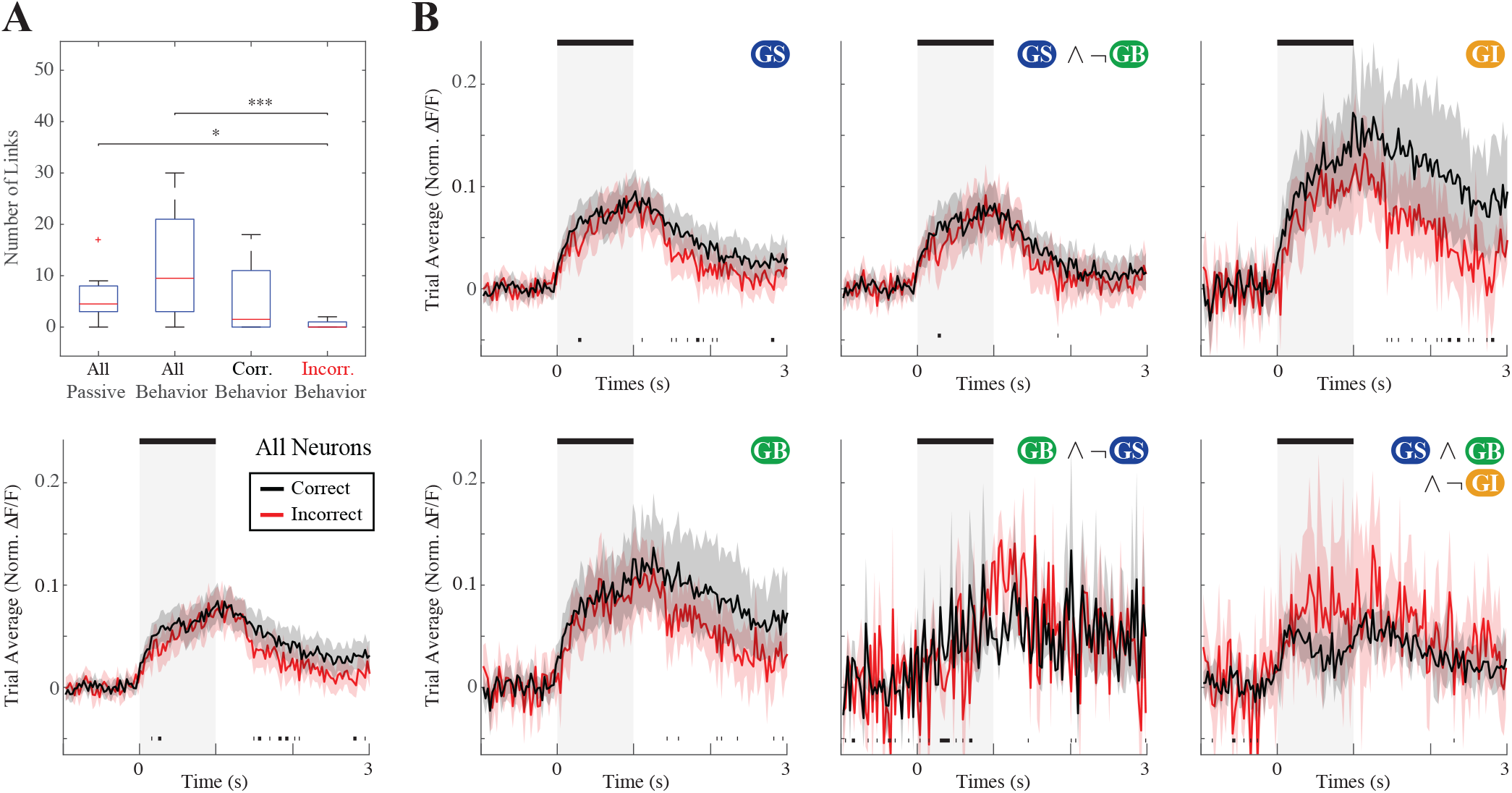
GC network and G-taxonomy analysis of mouse A1 during tone discrimination. A) number of GC links between neurons across multiple experiments, during passive and behavior conditions (top, * *p <* 0.05; ** *p <* 0.01; *** *p <* 0.001). Average activity of all neurons during correct vs. incorrect behavior are shown at the bottom (*N* = 100 neurons). B) Average activity of different neuron categories from the proposed G-taxonomy during correct vs. incorrect behavior. GI neurons show notable divergence between correct vs. incorrect conditions that is sustained throughout the post-stimulus interval, whereas the GS and GB neurons exhibit more transient differences. Shaded hulls represent 95% bootstrapped confidence intervals, and black bars below each trace indicate groups of time points that show significant difference between correct vs. incorrect trials (at a 95% confidence level).

To further tease apart the response differences shown in Fig. 5–A, we leverage our proposed G-taxonomy and refine the trial averages based on the Granger sensori-behavioral relevance of the neurons. Fig. 5–B shows the aforementioned refinement. The leftmost column in Fig. 5–B shows the trial averaged responses of GS (top) and GB (bottom) neurons. The GS neurons show significant differences in correct vs. incorrect trials both during the peri-stimulus and post-stimulus intervals, whereas the GB neurons show different responses later during the trial. Given that some GS neurons may also be GB and some GB neurons may also be GS, we further inspect the average responses of neurons that are GS and *not* GB (i.e., exclusively GS, top) and those that are GB and *not* GS (i.e., exclusively GB, bottom) in the middle column. Interestingly, the exclusively GS neurons show significant differences between correct vs. incorrect trials early during the stimulus presentation, which does not extend beyond the stimulus off-set and through the rest of the trial. The exclusively GB neurons, however, exhibit response differences mainly during the pre- and peri-stimulus intervals. These two results suggest that exclusively GS neurons show differences that are specific to the stimulus presentation window and does not persist throughout the trial, and exclusively GB neurons show differences early on and prior to stimulus presentation, which could relate to the behavioral bias of the animals independent of the stimulus category.

In the third column of Fig. 5–B, we inspect the responses of GI neurons (top), as well as GS and GB neurons that are *not* GI (bottom). The GI neurons exhibit significant response differences between the correct and incorrect trials, which persists throughout the entire post-stimulus interval. This is consistent with the findings of [20], showing the reverberation of intersection information during the trail. The GS & GB neurons which are not GI, however, show transient response differences mainly in the pre-stimulus interval, in stark contrasts to the differences exhibited by the GI neurons. This result suggests that being GS & GB is not sufficient for the to neuron carry sensori-behaviorally relevant information, whereas GI neurons carry information from both the sensory stimulus and behavior that persists throughout the trial.

Finally, we probed the response differences in incorrect vs. correct trials refined by the behavioral outcomes of lick or no-lick in Fig. 6. The top row shows the responses of GS (left), GB (middle), and GI (right) neurons during the lick trials, in which correct and incorrect outcomes correspond to H and F, respectively. The GS neurons show response differences shortly after the stimulus off-set, whereas the GB and GI neurons exhibit response differences that persist throughout the trial, more prominently for the hit trials. The bottom row similarly shows the responses during no-lick trials, in which correct and incorrect outcomes correspond to C and M, respectively. In contrast to the top panel, for the no-lick trials there are no notable differences between the responses of either category of neurons in C vs. M trials. This results suggests that the differences observed in Fig. 5 indeed depend on the animals’ actual response, and not just on the correct vs. incorrect categorization; while the H and F categories show significant response differences that are distinct for GS, GB, and GI neurons, the neuronal responses in M and C categories seem to be similar and not dependent on their functional type.

**Fig 6.**
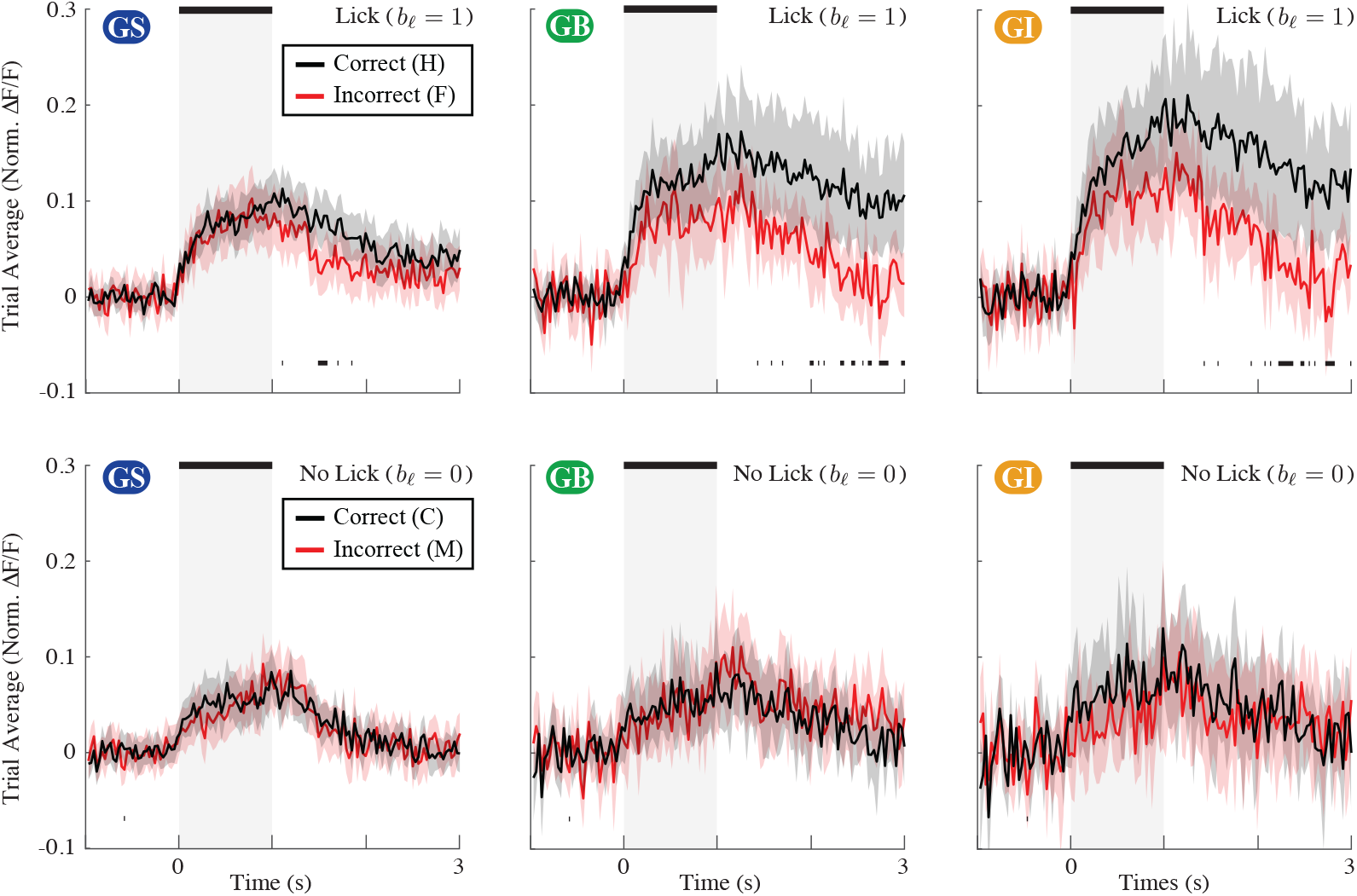
GC network and G-taxonomy analysis of mouse A1 during tone discrimination (cont’). Average activity of different neuron categories from the proposed G-taxonomy during correct vs. incorrect behavior for lick (top) and no-lick (bottom) trials. Consistent with Fig. 5, GI neurons exhibit sustained difference between correct and incorrect trials, but only for lick trials. During no-lick behavior, no significant difference in the activity of neurons across correct and incorrect trials is registered. Shaded hulls represent 95% bootstrapped confidence intervals, and black bars below each trace indicate groups of time points that show significant difference between correct vs. incorrect trials (at a 95% confidence level).

## Discussion

We introduced a unified statistical framework for joint estimation of GC influences among neurons in an ensemble as well as a functional taxonomy, namely G-taxonomy, that quantifies the sensori-behavioral relevance of each neuron, all *directly* from two-photon calcium imaging data. Our proposed G-taxonomy labels a neuron as: (1) Granger Stimulus (GS), implying that the stimulus is a significant predictor of its activity, (2) Granger Behavior (GB), implying that the neuron’s activity is a significant predictor of the behavior, and (3) Granger Intersection (GI), implying that the neuron encodes sensory information that is relevant to behavior. The GC links among neurons as well as the foregoing functonal G-taxonomy are established using rigorous statistical tests. Using both simulated and experimental recordings from mouse auditory cortex, we demonstrated the utility of the proposed framework in identifying the GC links between the neurons as well as establishing their sensori-behavioral relevance.

Our proposed approach is inspired by the Intersection Information (II) framework [15, 16, 20, 22, 51], in which the partial information decomposition (PID) machinery is used to quantify how much information, in the sense of Shannon, a neuron carries about the sensory stimulus, the behavioral readout, or both. While the II framework has successfully been applied to a wide variety of neural data and has resulted in novel insights regarding the interaction of sensory encoding and behavior, it comes with a number of challenges: First, due to the exponential complexity of the PID framework in the number of neurons, the II framework has mainly been restricted to a pair or triplets of neurons or variables; Second, the II framework assumes that the spiking activity is directly observed or its reliable estimates from indirect measurements are available; Third, the II framework represents the stimulus by simple features chosen *a priori*, often constant and univariate, in favor of the ease of solving the optimization problems involved in PID. As such, application of the II framework to larger networks of neurons, presented with potentially complex and time-varying stimuli, and measured indirectly via modalities such as two-photon imaging.

Our approach addresses these shortcomings through several mechanisms: First, by adopting the Granger machinery, as opposed to PID, we use a hierarchical parametric generative model that admits efficient estimation and scales favorably with network size. This is in contrast to the common applications of II that use nonparametric and model-free estimates of mutual information between pairs or triplets of neurons. Leveraging the milder dependence of estimation accuracy to the sample size in parametric models, we form a series of rigorous statistical tests that allow family-wise error correction and thus preserve statistical power. In the II framework, however, nonparametric tests are often used, which are known to have less statistical power than parametric ones, especially when the number of trials and their duration are limited.

Second, the parametric nature of our hierarchical models allows us to represent the stimuli as a multivariate time-varying signal, whose effects are parameterized by high-dimensional receptive fields. As such, no *a priori* assumptions are made on which features of the stimuli are relevant, and instead these features are captured in a data-driven fashion through the estimated receptive fields.

Finally, while most existing methods perform network inference in a two-stage fashion, by first estimating spikes and then forming network measures from said spike estimates, our modeling and estimation framework integrates several techniques such as dynamic Bayesian network models, state-space inference, Pólya-Gamma augmentation, and variational inference to avoid two-stage processing and thereby minimize estimation biases.

Another key advantage of our proposed framework is to *simultaneously* estimate the functional connectivity between the neurons in the sense of Granger, and to attribute a sensori-behavioral taxonomy to all the neurons, also in the sense of Granger, to quantify their role in the network function. Most existing methods solve these two inverse problems independently and disjointly: they either focus on capturing the encoding or readout properties of the neurons, regardless of their connectivity, or just aim at estimating functional connectivity by ignoring the effects of the stimulus. The advantage of solving these two problems simultaneously is two-fold: First, in detecting the GC links between neurons, the effect of external stimuli as a confound would be ruled out. For instance, if two neurons are being driven by the same stimulus with different latencies, by explicitly modeling the receptive fields and quantifying the stimulus effect, the detected GC links would only be resulting from the coupling of the endogenous latent process governing the neuron’s stimulus-independent activity; Second, in attributing the sensori-behavioral relevance to each neuron, the network effects arising from endogenous interactions between the neurons as a confound would be ruled out. For instance, we can rule out if heightened activity of a neuron corresponding to a particular behavioral category is due to its relevance to behavior, or is just due to receiving an endogenous input from another neuron that contributes to behavior.

We presented a carefully designed simulation study that showcases the intricacies involved in attributing sensori-behavioral relevance to neurons in a network and demonstrated that our proposed framework reliably achieves this task, while existing two-stage approaches suffer from poor hit rate and high false alarms.

We also applied our proposed framework to two datasets of two-photon recordings from mouse A1 layer 2/3. The first dataset corresponded to a network of mouse A1 neurons during passive listening in a sequence of trials in which either a broad-band white noise auditory stimulus is on or off in a randomly interleaved fashion [38]. Our proposed method labeled a set of neurons as GS neurons, which upon further inspection of their trial averaged activity, exhibited highly distinct responses in stimulus on vs. off trials. Interestingly, while some neurons exhibited distinct responses in these two conditions, they were not labeled as GS; indeed, the difference in their activity was much later throughout the trial in the late post-stimulus period.

The second dataset was collected during a tone discrimination task [50], in which the mice were trained to lick a spout after the onset of low frequency target tones and avoid licking when high frequency non-target tones are presented. Summarizing the GC network and G-taxonomy statistics across several animals resulted in several key findings: First, comparing the GC networks between the passive and behavioral conditions of correct vs. incorrect showed that the incorrect GC networks are sparser than the passive condition, consistent with the results of [50]. Second, neurons labeled as GI showed persistent differences in their responses between correct vs. incorrect trials throughout the post-stimulus period, also consistent with the reported reverberation of information in II neurons [22, 50].

Third, neurons that were exclusively labeled as GS only showed transient response differences between correct vs. incorrect trials in early peri-stimulus period, and those labeled as exclusively GB, showed transient response differences predominantly in the pre-stimulus period. This result suggest that these neurons may not play a key role in linking sensory stimulus to behavioral outcome as their activity profile does not correlate with the animal’s behavioral category in a persistent fashion. Finally, neurons that were labeled as GI showed a striking difference in their activity profile, persistently during the post-stimulus period. Interestingly, the activity of neurons labeled as GS and GB, but not GI, as starkly contrasting to those labeled as GI. This result is consistent with the rationale of the II framework, implying that neurons that encode sensory stimulus and are predictive of behavior, may not necessarily encode features of the stimuli that are relevant to behavior.

While our proposed framework successfully recovers GC links and sensori-behavioral relevance directly from two-photon imaging data, several limitations remain. The current model assumes linear vector autoregressive dynamics and fixed time lags between neural influences, which may not fully capture nonlinear or time-varying dependencies present in real cortical circuits. Future work could incorporate kernelized or hierarchical extensions to better represent complex neuronal response properties. Additionally, analyses were performed on relatively small neuronal subsets due to computational constraints. Extending this framework to full-field datasets with hundreds of neurons will require optimized inference and regularization strategies. Finally, although GC reflects predictive influence, it does not imply direct synaptic connectivity. Combining this approach with anatomical tracing or targeted optogenetic perturbations will be essential for establishing causal validation.

In summary, this study presents a unified and statistically grounded framework for uncovering directed functional interactions and sensori-behavioral relevance of network activity in the sense of Granger from two-photon data. By jointly modeling stimuli, neural activity, and behavioral responses, it captures how sensory information can dynamically reorganizes network connectivity across behavioral states. Beyond auditory processing, the same principles can be extended to other sensory systems and experimental paradigms, offering a powerful tool for studying how distributed neural populations transform sensory input into behaviorally relevant computation.

## Methods

We first describe the generative model in detail, followed by the inverse problem and the proposed solution. The proposed inverse solution consists of three main parts: 1) efficient network parameter estimation, 2) full and reduced models used in the Granger formalism, and 3) statistcal testing of GC links and GS, GB, and GI attributes.

### Generative Model Formulation

Consider a standard experimental setting where we have the fluorescence traces of *N* neurons within a trial of length *T* frames for a total of *L* independent trials. We 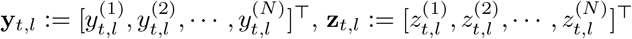, and 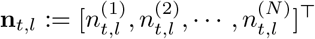 as the vectors representing the noisy observations, intracellular calcium concentrations, and ensemble spiking activities, respectively, at trial *l* and frame *t*. We aim to capture the temporal patterns within **y**_*t,l*_ using the following state-space model [38, 40, 52]:

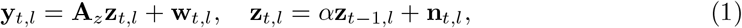

where **A**_*z*_ ∈ ℝ^*N* ×*N*^ represents a diagonal scaling matrix capturing the cross-neuron intensity variations, **w**_*t,l*_ is zero-mean i.i.d. Gaussian noise with covariance **Σ**_*w*_, and 0 ≤ *α <* 1 is the state transition parameter capturing the calcium decay dynamics via first-order autoregression. Note that our chosen state-space model exhibits non-Gaussian characteristics due to the binary nature of the spiking activity, that is, 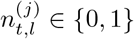

To model the spiking data, we employ two covariates within a point process or Generalized Linear Model framework with Bernoulli statistics. The first covariate, designated as the latent endogenous process 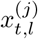, accounts for spontaneous and stimulus-independent neural activity. The second component models the responses driven by external stimuli **s**_*t,l*_ ∈ ↦^*J*^, where *J* denotes the flattened length of a multi-variate stimulus and its previous lags effective at time *t*. The latter component is denoted by the exogenous component. We hereafter assume that the latent endogenous process evolve over a comparatively slower time scale relative to the imaging frame rate, i.e., observations within a window of length *W* ≥ 1 are driven by the same instances of the latent process [38]:

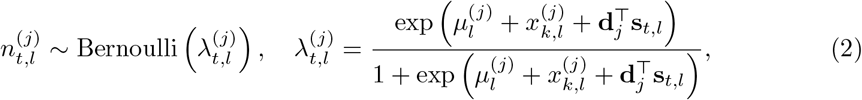

where 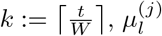 is the baseline rate parameter of neuron *j* in trial *l*, and **d**_*j*_ is the flattened receptive field of neuron *j* encoding the effect of stimulus **s**_*t,l*_. The non-linear mapping relating the different covariates to 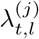 is referred to as the logistic link, which is the canonical link for a Bernoulli process within the Generalized Linear Model framework [53, 54].

To account for the endogeous and directed coupling of the neurons, we employ a vector autoregressive (VAR) process to model the dynamics of 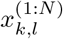. Letting 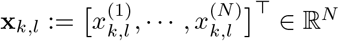, the VAR model can be expressed as follows [55]:

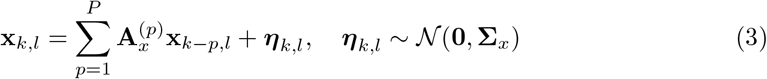

where 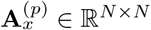 represents the AR coefficients associated with the *p*^th^ time lag, and the noise covariance **Σ**_*x*_ ∈ ℝ^*N* ×*N*^ is a diagonal matrix with the *j*^th^ diagonal entry given by 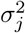.

Finally, we use a categorical probabilistic model to account for observed behavior.

Letting 𝒯_RO_ ⊆ *{*1, 2, · · ·, *T}* be the readout interval (e.g., the second half of the stimulus interval), the binary behavior *b*_*l*_ (e.g., lick/no-lick) in trial *l* is modeled as

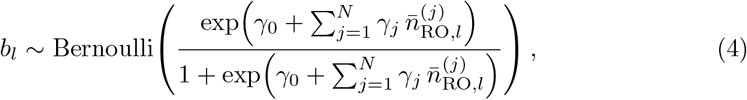

where

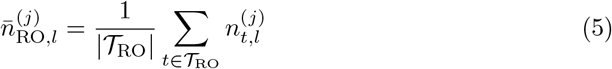

is the average activity of neuron *j* in trial *l* during the readout interval 𝒯_RO_, *γ*_0_ is the behavior bias parameter, and *γ*_*j*_ capture the contribution of the different neurons to the behavior. Fig. 1 illustrates the key elements of the foregoing generative model.

The overall inverse problem can be stated as follows: given the observed fluorescence traces from *N* neurons over *L* independent trials of duration *T* each, 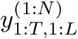, extract the Granger Causality (GC) links among the neurons as well as the proposed G-taxonomy of GS, GB, and GI for all the *N* neurons in the population.

The objective of the inverse problem hinges on accurately estimating the network parameters 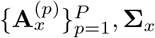, the receptive fields 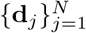, and behavior parameters 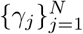. Then, to test whether a neuron *i* has a GC influence on neuron *j*, we first estimate the network parameters without imposing any restrictions, namely the *full* model. Then, we set the coupling VAR parameters from neuron *i* to neuron *j* in the and re-estimate all the other network parameters, resulting in the *reduced* model. If including 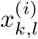 in the prediction of 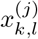 results in significantly higher likelihood, the *full* model is preferred over the *reduced* model, indicating the existence of a GC link from neuron *i* to *j* [34]. A similar procedure can be applied to the inference of G-taxonomy, albeit with more technical care in defining the respective *full* and *reduced* models.

In the subsequent sections, we first give a detailed account of the network parameter estimation procedure, followed by defining the *full* and *reduced* models for G-taxonomy and the associated statistical tests.

### Inverse Solution (1): Efficient Network Parameter Estimation

Let 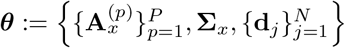 be the set of network parameters to be estimated. To this end, we use the EM framework, which offers an iterative procedure for obtaining a sequence of parameter estimates 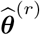 such that:

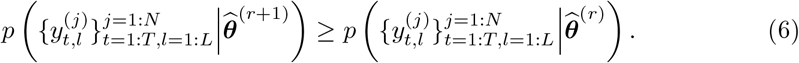

The overall architecture of the EM procedure used for parameter estimation in our work is shown in Fig. 7. We illustrate this procedure for the *full* model first. The joint log-likelihood of the observations, latent processes and parameters can be expressed as:

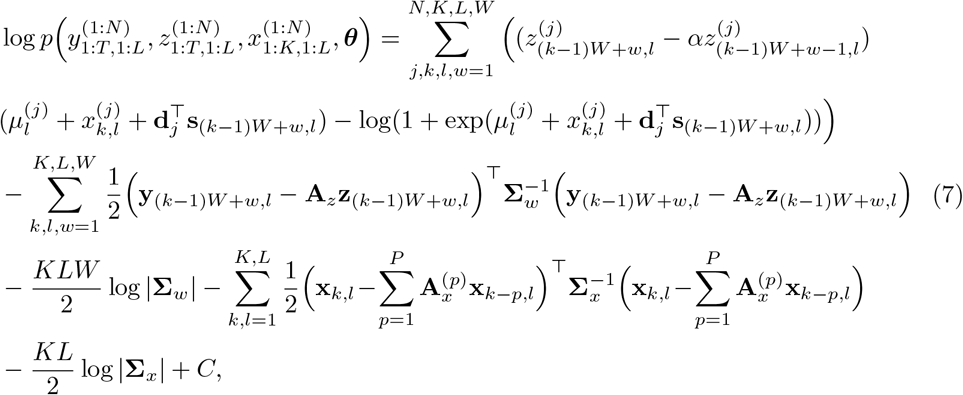

where *C* stands for terms that are not functions of 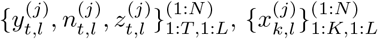 or ***θ***. Hereafter, we use shortened notations 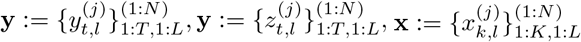, and 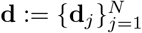 for convenience.

**Fig 7.**
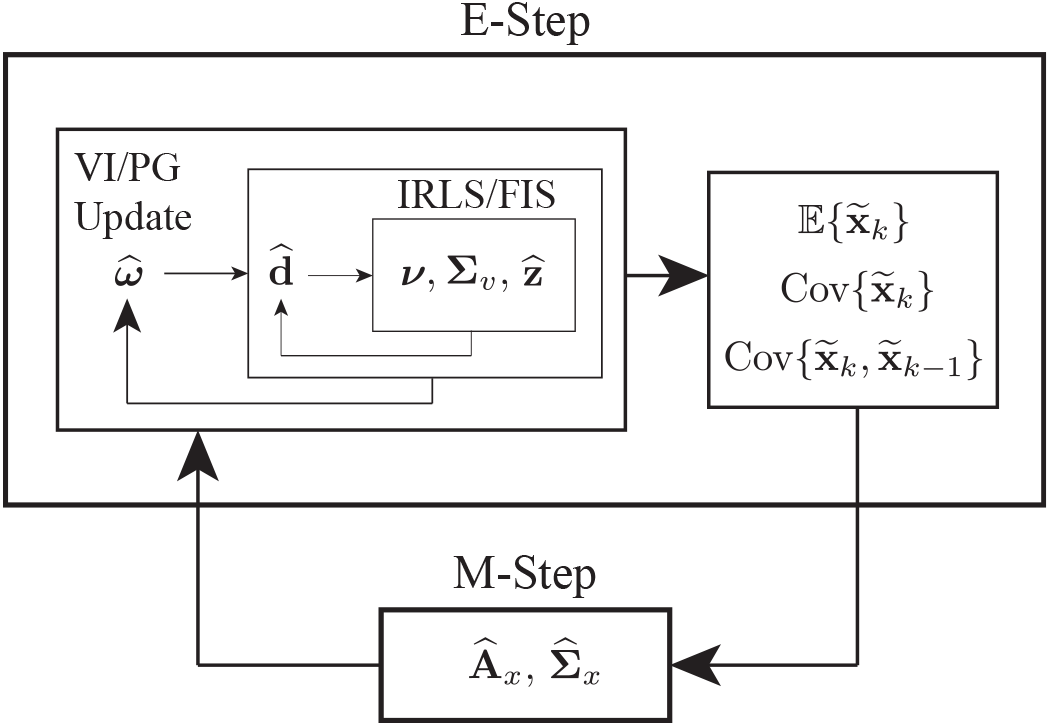
Schematic depiction of the inverse solution. The E-step includes two nested iterative procedures, namely variational inference via Pólya-Gamma augmentation (VI/PG) and Iteratively Re-weighted Least Squares/Fixed Interval Smoothing (IRLS/FIS). The first and second moments of the latent endogenous process are then computed and fed to the M-step to compute the network parameters.

### The E-Step

We need to compute the so-called Q-function given by:

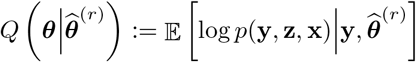

where 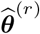 is the parameter estimates at iteration *r*. Utilizing the state augmentation technique, we express the VAR model as:

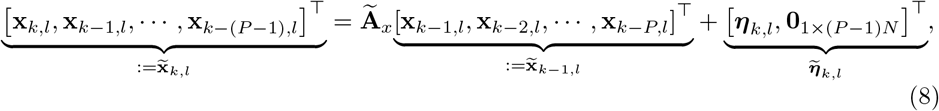

with

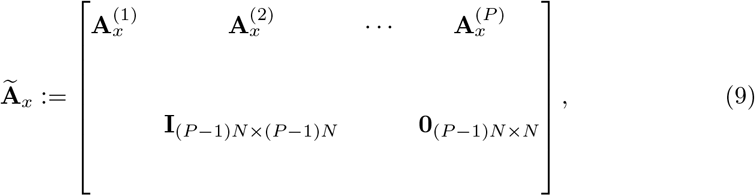

and 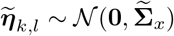 where

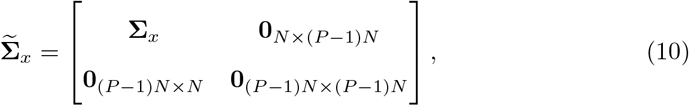

complete log-likelihood of Eq. (7) can be more concisely expressed as:

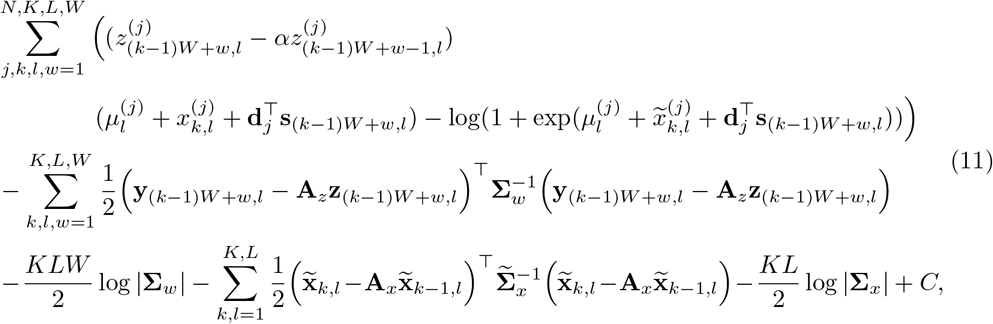

Upon inspecting Eq. (11), the complexity of the E-step becomes evident: the joint log-likelihood function involves interactions that are not purely quadratic, hence the expectations can not be expressed using the first and second moments, as in the Gaussian case. To address this, we use an instance of Variational Inference (VI) [56–58], a powerful technique from Bayesian statistics. VI approximates complex posterior distributions by transforming the inference problem into an optimization problem. This method reduces computational complexity compared to traditional approaches based on sampling, providing an efficient and scalable solution for parameter estimation.

To proceed with the VI framework, we assume that the constants *α*, **A**_*z*_ and **Σ**_*w*_ are either predetermined or can be reliably estimated from pilot trials. For instance, we treat **A**_*z*_ as a diagonal matrix, estimating its diagonal entries from the intensity of spiking events. Moreover, *α* can be estimated based on the decay rate of observed spikes, as it relates to the calcium indicator and imaging technique. Finally, **Σ**_*w*_ can be estimated using the background fluorescence in intervals without spiking activity.

### Variational Inference: Decoupling via Pólya-Gamma Augmentation

When confronted with problems encompassing both discrete and continuous random variables, the straightforward utilization of VI frequently results in the generation of intractable probability density functions. In our specific model, deriving an appropriate variational distribution for **x**_*t,l*_ using standard distributions becomes intractable due to the interrelationships between Bernoulli and Gaussian random variables in the posterior. To address this challenge, we utilize the Pólya-Gamma (PG) latent variable augmentation technique [41, 59, 60]. By examining Eq. (7), we observe that the posterior density, *p*(**x**|**z, x, A**_*x*_, **d, Σ**_*x*_), admits conditional independence in *k* and *l*, through expressing:

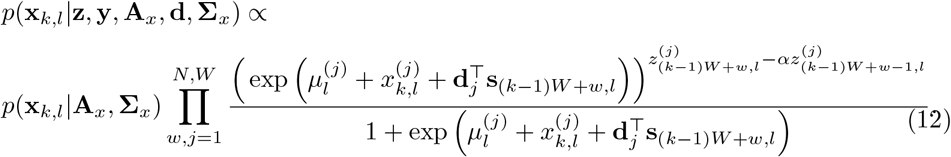

This latter density possess the desired characteristics for the PG augmentation scheme which relies on the following identity [38, 41]:

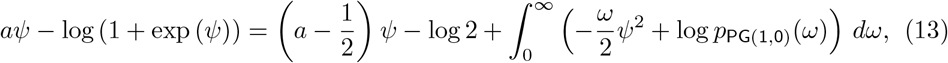

where *p*_PG(1,0)_(*ω*) represents the PG density. Comparing the first term in Eq. (11) with the left-hand side of Eq. (13), we can identify

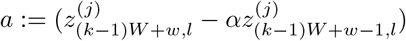 and 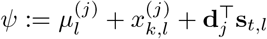. We then introduce a set of i.i.d. auxiliary latent random variables 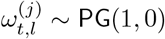 for 1 ≤ *j* ≤ *N*, 1 ≤ *t* ≤ *T* and 1 ≤ *l* ≤ *L*. Letting 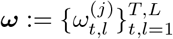, we have:

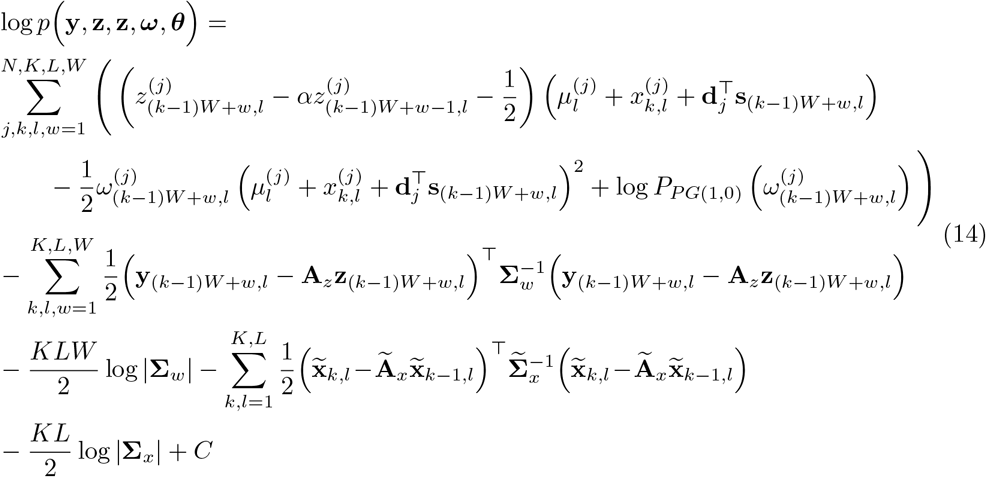

By taking 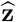 and 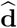 as parameters and assuming their estimates are available, we next assume a mean-field variational density family and adopt the Coordinate Ascent Variational Inference (CAVI) procedure to derive the optimal variational densities [38]. The procedures for updating the parameters 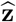 and 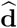 will be detailed in the subsequent section.

The augmented log-likelihood in Eq. (14) exhibits a *quadratic*dependence on 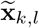 and **z**_*t,l*_. Consequently, the conditional distribution 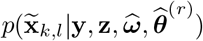, given an estimate 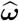, can be identified as Gaussian. This property enables us to leverage fixed interval smoothing [42] and covariance smoothing [46] for efficient computation of the conditional expectation for 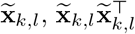, and 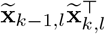 given 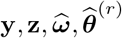, to fully characterize the expectations in the E-step.

The augmented log-likelihood in Eq. (14) also implies the conditional distribution 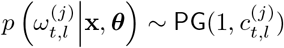, with 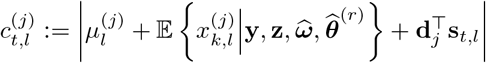, where *t* := (*k* ™ 1)*W* + *w*. Therefore, we estimate the PG variables as 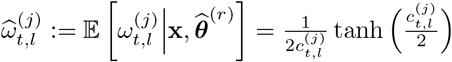. To improve the accuracy of the latent PG variable estimation, we alternate between estimating the first and second moments of 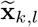 and 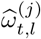, before proceeding to the M-step.

### VI Parameter Updates

Recall that we treat **z** and **d** as an unknown parameters, instead of unknown random variables whose densities would require variational parametrization. This choice is due to the temporal dependencies induced by the interaction of **z** and **x** in Eq. (7, which complicate quantifying the mean-field variational densities.

Note that the variable 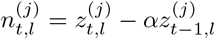 is a Bernoulli random variable, posing a set of constraints 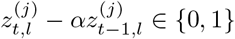, for *t* = 1, · · ·, *T, l* = 1, 2, · · ·, *L*, and *j* = 1, · · ·, *N*. The combinatorial nature these constraint renders the estimation of **z** intractable. Following the relaxation procedure of [38, 43], we instead use the modified log-likelihood:

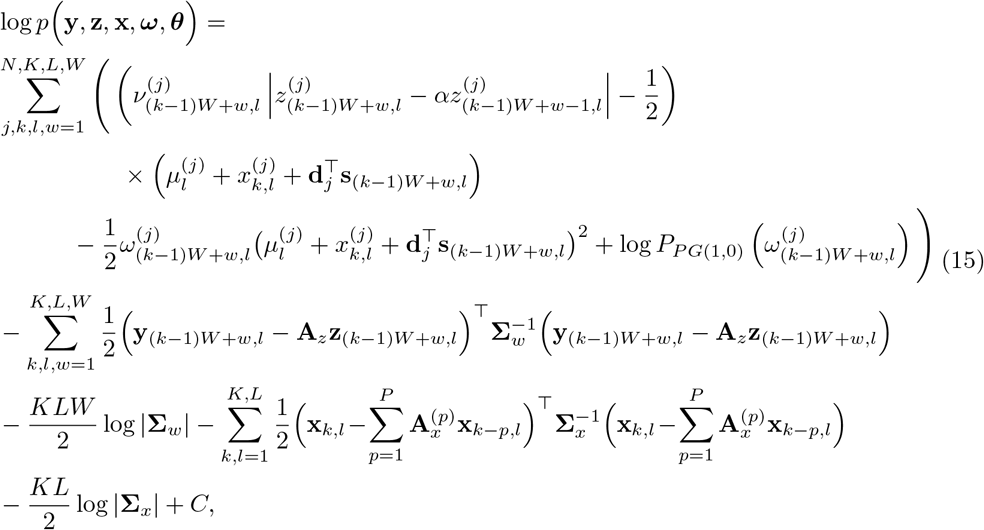

where 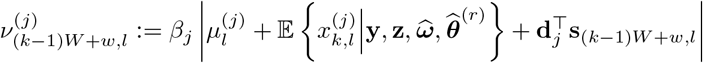 with *β*_j_ ≥ 0 denoting a hyper-parameter. In our analysis, we set the value of *β*_*j*_ inversely proportional to the variance of the observed fluorescence signal as *β*_*j*_ = 0.7*/*(**Σ**_*w*_)_*j*_. To maximize the modified log-likelihood in Eq. (15) with respect to **z**, we utilize the Iteratively Re-weighted Least Squares (IRLS) procedure [44]. At iteration *i* given a current estimate 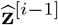, the log-likelihood can be identified as that of the state-space model:

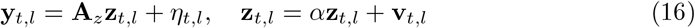

In this equation, *η*_*t,l*_ ~ 𝒩 (0, **Σ**_*η*_) and 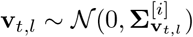 where 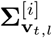 is a diagonal matrix with 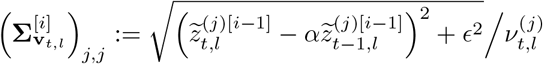, for some small constant *ϵ >* 0. Thus, the next iterate 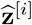 can be efficiently computed using using FIS [42]. Note that this method for estimating the calcium concentrations **z**_*t,l*_ functions as a *soft* spike deconvolution, seamlessly integrated within the variational inference framework. This differs significantly from traditional two-stage estimators, which rely on *hard* spike deconvolution.

With the updated estimates of 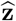, the kernel parameters **d** can be estimated by maximizing the log-likelihood in Eq. 14 with respect to **d**_*j*_:

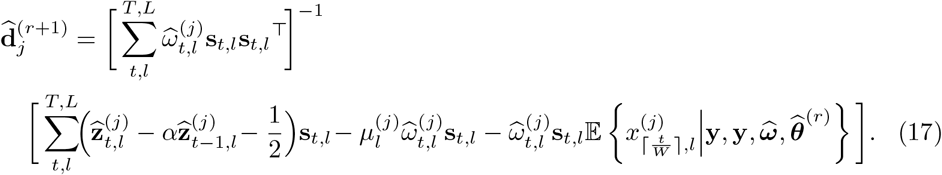

Finally, yhe VI procedure iterates between updating the variational densities and the parameters.

### The M-Step

In the M-step the parameters **A**_*x*_ and **Σ**_*x*_ are updated to maximize the Q-function. The update rules for the *j*th row of **A**_*x*_ and **Σ**_*x*_ are given by:

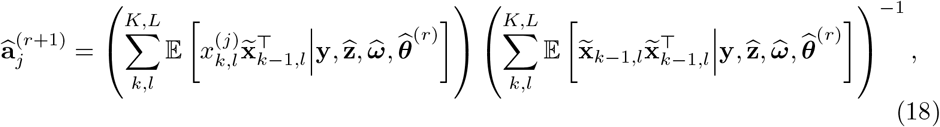

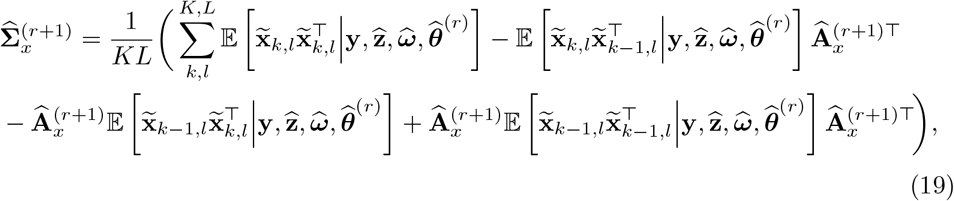

The EM algorithm proceeds until a standard stopping criterion is met [38].

### Inverse Solution (2): Full and Reduced Models in the Granger Formalism

Given the foregoing efficient procedure to estimate the network parameters, we next need to define the nested *full* and *reduced* models for the GC links as well as the G-taxonomy to be able to extract relevant statistics for hypothesis testing.

### Full and Reduced Models for GC Links Between Neurons

In the reduced model for assessing a GC link from neuron *i* to neuron *m*, we remove the influence of the source neuron on the target neuron in Eq. (3) by setting the corresponding VAR coefficients to zero. The update rule of Eq. (18) for all *j*≠ *m* remains the same. For *j* = *m*, however, we utilize a reduced vector 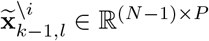 derived by removing the elements of 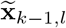 that correspond to neuron *i*. The update for 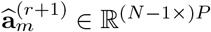 is then given by:

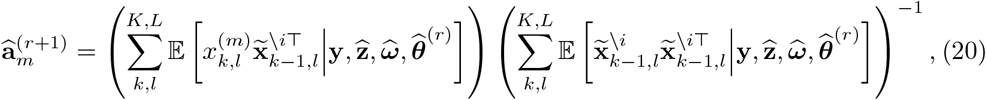

The coefficients corresponding to neuron *i* are then explicitly set to zero. Similarly, Eq. (19) is modified by replacing 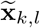 and 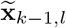 with 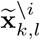 and 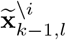, respectively.

### Full and Reduced Models for GS and GB Neurons

In the reduced model for assessing the effect of the stimulus on neuron *m*, we set **d**_*m*_ = **0** and estimate all the other network parameters. In other words, Eq. (2) for all *j*≠ *m* remains the same, and we have:

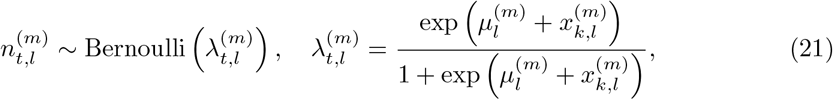

The reduced model for assessing the effect of the activity of neuron *m* to behavior, corresponds to setting *γ*_*m*_ = 0, i.e.:

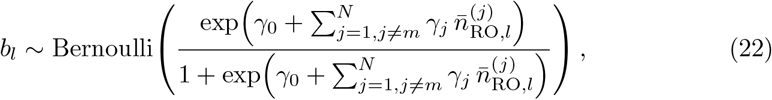

The parameters *γ*_*j*_, *j*≠ *m* can be estimated by maximizing the joint log-likelihood of 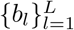 using the Newton–Raphson algorithm.

### Full and Reduced Models for GI Neurons

Recall that a neuron *m* being both GS and GB does not imply that it encodes stimulus information which is subsequently utilized for task-related behavior, i.e., a GI neuron. Mirroring the rationale of II theory based on mutual information, here we need to see if estimating the stimulus from the average activity of neuron *m* during the readout window would be significantly improved if the behavior is included as a covariate or not. Noting that the behavior is typically a categorical variable and the average activity of neuron *m* during the readout window is constant, using them to estimate the time-course of the multi-dimensional stimulus does not constitute a rich enough model to test the aforementioned effect, since both the full and any possible reduced model are likely to be equally and drastically poor. We thus suppress the temporal dimension of the stimulus **s**_*t,l*_, and consider a categorical representation for *s*_*l*_ ∈ *{*1, 2, · · ·, Ξ*}*. For instance, if the stimulus corresponds to a modulated presentation of different pure tones, the Ξ categories correspond to the number of tones to generate the stimulus set.

To this end, we use a multinomial logistic model as the *full* model:

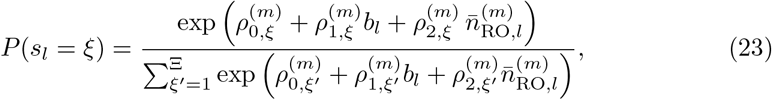

where the parameters 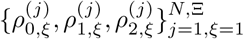 are estimated by maximizing the multinomial joint log-likelihood of 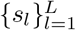 using the Newton–Raphson algorithm.

The reduced model can then be formulated by constraining 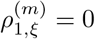, thereby removing the direct influence of the behavior on the predicted stimulus category. As such, the reduced model isolates the contribution of the activity of neuron *m* during the readout interval to stimulus prediction, independent of the behavior. The rest of the parameters 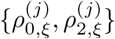 are re-estimated under the constraint 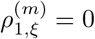 using the same optimization procedure.

### Inverse Solution (3): Statistical Testing of GC Links and GS, GB, and GI Attributes

The test statistic to be used to compare the full and reduced models is the deviance difference [34, 35]:

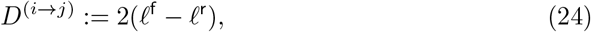

where *ℓ*^f^ and *ℓ*^r^ denote the log-likelihood of the observations corresponding to the full and reduced models, respectively. Given that the latent endogenous process is unobserved, computing the log-likelihood of the observed fluorescence traces is not straightforward, as it involves intractable integrations over the range of the latent process. We thus adopt the procedure of [61] that uses a specific sample path of the latent endogenous process that significantly simplifies the computation of the log-likelihood. The key idea is to factorize the likelihood of the observations as:

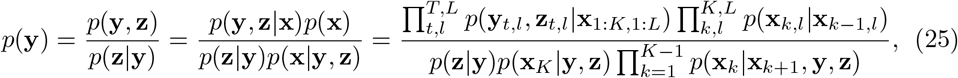

where **x** and **z** can be taken as *any* possible choice of the latent endogenous process and calcium concentrations. The numerator of Eq. (25) can be expressed in closed-form using our generative model. However, the denominator lacks such simplicity, primarily due to the intricate interplay between **x**_*k,l*_ and the observation model. To address this complexity, we employ the Laplace approximation by making the assumption 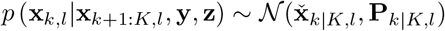, where 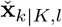 and **P**_*k*|*K,l*_ satisfy [61, 62]:

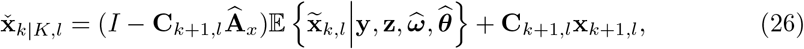

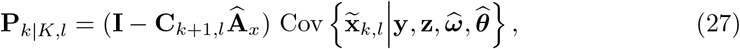

where

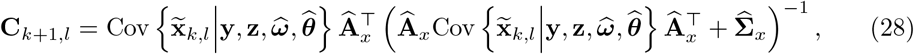

with 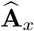 and 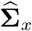 being the estimated AR coefficients and the first and second moments of 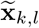 being available following the parameter estimation procedure. Initializing with 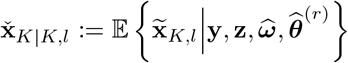, for *k* = *K* − 1, *K* − 2, · · ·, 1, we proceed in a backward fashion by setting 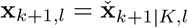 in Eq. (26) to obtain a sample path 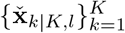. Evaluating the denomintator of Eq. (25) at this sample path yields 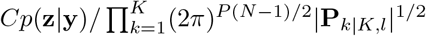. Finally, we evaluate the likelihood at the estimated 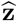 from the parameter estimation procedure. The simplified log-likelihood of the observations can be express as:

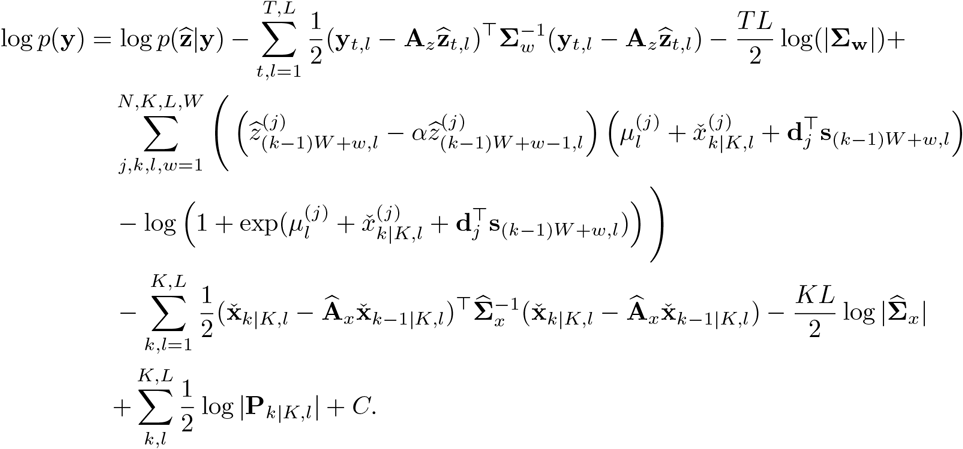

Note that in the absence of observation noise **w**_*t,l*_, we have **y**_*t,l*_ = **A**_*z*_**z**_*t,l*_, and therefore 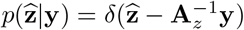. Assuming that the observation noise is small, and the prior *p*(**z**_*t,l*_) induced by the VAR process **x**_*t,l*_ is smooth around 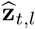, the posterior 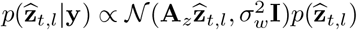 is approximately Gaussian with mean 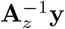 and covariance 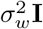 and is therefore negligibly dependent on the network parameters.

Following this approximation, 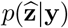 takes similar values in the full and reduced models. Therefore, log 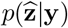 cancels out in computing *D*^(*i* ↦*j*)^. In fact, in order to speed up the computation of the estimation framework, one can use the regularization parameters 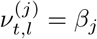 independent of the network parameters, so that 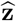 can be computed once for the full model only and used in the reduced model. This way, 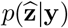 in the full and reduced model is evaluated at the same estimate 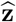.

The foregoing expression of the log-likelihood of the observations can be used to compute *D*^(*i*↦*j*)^ as well as *D*^(*S*↦*j*)^, denoting the deviance differences of the effect of neuron *i* to *j* and the stimulus to neuron *j*, respectively. The log-likelihood of the full and reduced models for the behavior can be computed using the closed-form expression of the log-likelihood of Bernoulli variables, resulting in the deviance difference *D*^(*j* ↦*B*)^. Same goes for the log-likelihood of the stimulus categories in the full and reduced models corresponding to GI neurons, which provides *D*^(*B* ↦*S*|*j*)^.

Asymmptotic analysis of the deviance difference *D*^(*i* ↦*j*)^ involves the properties of the central and non-central chi-squared distributions [34, 35, 63–65]. Let *M* be the number of parameters that are removed in the reduced model. Then, in the absence of a GC effect, *D*^(*i* ↦*j*)^ asymptotically (in the observation length) follows a chi-squared distribution with *M* degrees of freedom *D*^(*i* ↦*j*)^ ~ *χ*^2^(*M*), and in the presence of a GC effect, it follows a non-central chi-squared distribution, *χ*^2^(*M, ν*^(*i* ↦*j*)^) with the same degree of freedom *M* and non-centrality parameter *ν*^(*i* ↦*j*)^. The non-centrality parameter *ν*^(*i*,↦*j*)^ can be estimated as 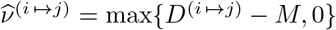 [34]. The degree of freedom *M* for assessing the different effects in our framework are given as:

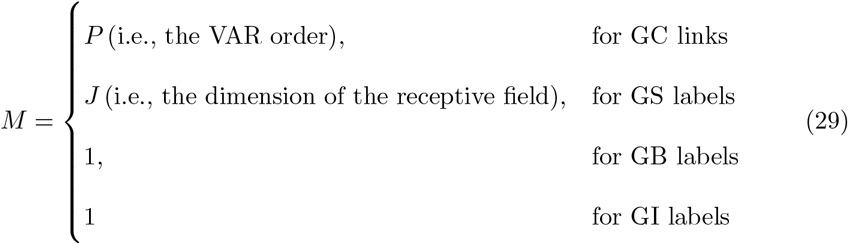

The p-value of the test comparing the full and reduced model based on *D*^(*i* ↦*j*)^ is therefore computed as 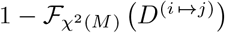, where 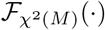 is the CDF of *χ*^2^(*M*). For a target significance level of *δ*, if we detect a GC link using the thresholding rule 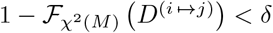, we can control the false discovery rate (FDR) across all estimated GC link by applying the Benjamini–Yekutielli (BY) procedure [48], at an average FDR of 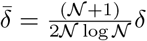, where 𝒩 is the number of simultaneous tests.

To summarize each test, we utilize the J-statistic associated with the link *i* ↦ *j*, defined as

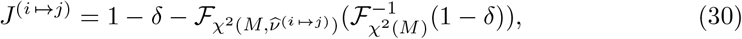

with 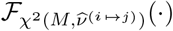 denoting the CDF of and 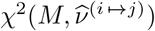. Values of *J*^(*i*↦*j*)^ ≈ 0 indicate small evidence to reject the null hypothesis, whereas *J*^(*i*↦*j*)^ ≈ 1 denote strong support for the alternative hypothesis. Thus, the J-statistic, bounded within [0, 1], serves as an interpretable index of the strength associated with each inferred GC effect. Similarly, we can assign the following J-statistics to the detected GS and GB neurons:

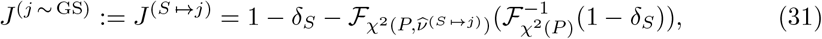

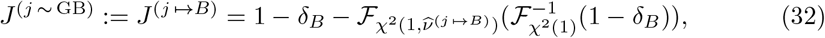

where *δ*_*S*_ and *δ*_*B*_ denote the target significance levels for the GS and GB categories, respectively, and are determined given a target FDR level and the number of simultaneous tests.

For a GI neuron *j*, we define 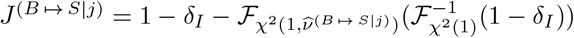, where *δ*_*I*_ is the target significant level for the GI category. Given that three difference statistical tests are required to identify GI neurons, by the union bound, the sum of type-I and type-II error probabilities of attributing a GI label are bounded by

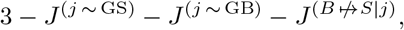

where *J*^(*B* ↛ *S*|*j*)^ := 1 − *J*^(*B* ↦ *S*|*j*)^. Subtracting this expression from unity yields the following lower bound on the J-statistic for a GI neuron:

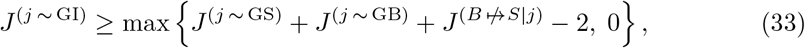

and can be used as measure of statistical confidence in labeling a neuron as GI.

### Statistical Testing of Trial Average Differences

In Figs. 5 and 6, for each sub-panel we computed the average difference between the correct and incorrect trials and obtained the point-wise 95% confidence intervals using bootstrapping. Then, we first identified individual time points with significant positive difference at a level of 5% (bootstrap test, one-sided). Among these individually significant points, starting from the most significant peak level, we iteratively lowered the level and included all the sub-level points as a group and recomputed the group-level *p*-value via Monte Carlo resampling (10, 000 surrogates), until the significance level crossed 5%. The final such group indicated the time points that show simultaneously significant positive differences. In the case of finding more than one peak at the same level, we performed Bonferroni correction for multiple groups to maintain the overall significance level at 5%. This procedure is adapted from [39, 66], which we refer the reader to for more details.

## Data Availability

Data and software implementations in MATLAB v2024b of the algorithms discussed are available at: https://doi.org/10.13016/ivfi-ftgv

## Acknowledgments

This work has been supported in part by the National Science Foundation Awards No. ECCS2032649 and OISE2020624 (to BB) and the National Institutes of Health Awards No. RO1DC017785 (to POK) and U19NS107464 (to BB and POK).

## Notes

https://doi.org/10.13016/ivfi-ftgv

### Competing Interest Statement

The authors have declared no competing interest.

